# Predictors of colony extinction vary by habitat type in social spiders

**DOI:** 10.1101/612432

**Authors:** Brendan L. McEwen, James L. L. Lichtenstein, David N. Fisher, Colin M. Wright, Greg T. Chism, Noa Pinter-Wollman, Jonathan N. Pruitt

## Abstract

Many animal societies are susceptible to mass mortality events and collapse. Elucidating how environmental pressures determine patterns of collapse is key for our understanding of social evolution. Using the social spider *Stegodyphus dumicola* we investigated the environmental drivers of colony extinction along two precipitation gradients across southern Africa, using the Namib and Kalahari deserts versus wetter savanna habitats to the north and east. We deployed experimental colonies (n = 242) along two 800km transects and returned to assess colony success in the field after two months. Specifically, we noted colony extinction events after the two-month duration and collected environmental data on the correlates of those extinction events (e.g., evidence of ant attacks, # prey captured). We found that colony extinction events at desert sites were more frequently associated with attacks by predatory ants as compared to savanna sites, while colony extinctions in wetter savannas sites were more tightly associated with fungal outbreaks. Our findings support the hypothesis that environments vary in the selection pressures that they impose on social organisms, which may explain why different social phenotypes are often favored in each habitat.

## INTRODUCTION

The evolution of sociality changes the ways in which species interact with their environments. Living in groups can confer the ability to construct larger refuges, subdue more profitable prey, mount amplified defenses against would-be enemies, and increase fitness through alloparental care (Krause & Ruxton 2002, Lubin & Bilde 2007). However, sociality is not without costs. Group-living organisms may suffer enhanced resource competition as they struggle to support a larger number of co-occurring conspecifics (Yip 2008, Mayer et al. 2018). They may become more conspicuous to predators (Wrona & Dixon 1991) or be more susceptible to disease (Côté & Poulin, 1991). These costs can accumulate to a critical level at which they outweigh the benefits conferred by sociality, causing a group to collapse or go extinct (Avilés and Tufino 1998).

Site-specific selection, through which discrete areas impose divergent selection on a species, is a relatively common feature in nature that often results in phenotypic diversity (Siepielski et al. 2009, Siepielski 2013, Caruso et al. 2017). For example, spatial and temporal variation in precipitation patterns (Siepielski et al. 2017), prey community structure (Drummond & Burghardt 1983), or interference competition (Riechert 1993) can all generate contrasting patterns of selection on individuals, often resulting in local adaptation. We propose that, like selection pressures predicting the survival or demise of individuals, pressures driving group extinction events might differ according to the environments in which a group resides.

Social spiders provide a convenient means with which to observe the relationship between the environment and group extinctions. Most species of spider are solitary and intolerant of conspecifics, however, social species are less aggressive overall and have higher conspecific tolerance than their solitary counterparts (Lubin & Bilde 2007, Harwood & Avilés 2018). This distinction allows social spiders to coexist in a communal web and work cooperatively on collective tasks, such as foraging (Whitehouse & Lubin 2005), web construction and maintenance (Purcell & Avilés 2008), alloparentental care (Whitehouse & Jackson 1998), and defense against predator attacks (Henschel 1998, Purcell & Avilés 2008, Yip & Rayor 2011, Wright et al. 2016). Social spiders also experience high rates of colony extinction (Aviles 1986, Henschel 1998).

Several hypotheses have been proposed to explain the high incidence of colony extinction events in social spiders. First, the transition to sociality in spiders is met with a transition from outbreeding to inbreeding, which results in low effective population sizes and a reduced ability to response to changing environmental pressures (Agnarsson et al. 2006, Settepani et al. 2017). Second, the evolution of sociality is associated with a decrease in cross-contextual aggressiveness and increased conspecific densities that renders social spiders susceptible to a diversity of ecological pressures (Pruitt et al. 2012). These pressures include invasion by foreign species of spiders (Cangialosi 1990, Pruitt 2012), increased detection rate by ant predators (Purcell & Aviles 2008, Keiser et al. 2015), increased susceptibility to starvation and local resource competition (Aviles et al. 2002, Yip 2008, Majer et al. 2018), and an increased propensity to spread socially-transmitted microbes (Henschel 1998, Keiser et al. 2018). Different habitats may vary in the severity of these pressures based on local conditions (e.g. prey availability, predator abundance). In this study, we evaluate whether environments with contrasting precipitation regimes differ in the factors associated with group extinction events. Precipitation has been identified as a key driver of selection across many systems (Purcell 2011, Hoffman & Avilés 2017, Siepielski et al. 2017) and, as we outline below, is a candidate variable that may influence the threats to the survival of social spider groups.

The social spider *Stegodyphus dumicola* (Araneae, Erasidae) occupies both arid and wet regions throughout southwestern Africa in colonies ranging from 1-2,000 spiders (reviewed in Aviles & Guevara 2017). *Stegodyphus dumicola* colonies in the wild experience mixed success. Though the species is reasonably common, colony-wide mortality rates can approach 90% in some habitats (Henschel 1998). The causes of these high mortality rates are potentially manifold, such as a failure to capture sufficient prey (Majer et al. 2018), succumbing to colony attacks performed by ants (Henschel 1998, Keiser et al. 2015), and the spread of fungus through the colony web (Henschel 1998). The high rates of colony extinction in *S. dumicola* renders this species as a convenient model with which to explore associations between habitat type and the correlates of colony extinction. Prior studies on *S. dumicola* have found that the relationship between colony traits and success differs predictably across wet vs. arid sites (Pruitt et al. 2018), hinting that the pressures driving colony success and collapse may differ across environments.

Here we evaluate the following *a priori* predictions: we predict that colonies at wet savanna sites will capture more prey than colonies at arid sites, because insect abundance and biomass increases in habitats with higher precipitation (Janzen and Schoener, 1968). However, increased insect prey parts within spider webs combined with higher moisture at wet sites may facilitate the growth of fungus. Thus, we predict colony extinctions will be more frequently associated with fungal outbreaks at wet sites than at arid sites. Prior studies at arid sites have demonstrated that predatory ants are a lethal and disruptive force upon *S. dumicola* (Henschel 1998, Keiser et al. 2015, Wright et al. 2017). Although the threat of predatory ants towards *S. dumicola* has not been directly assessed in wet habitats, increased ant abundance in habitats with greater primary productivity suggests that these pressures may be even stronger at wet sites (Kaspari et al. 2000). Accordingly, we predict that higher ant populations at wet sites will increase frequency of attacks by predatory ants, as well as colony mortalities due to ant attack. To evaluate these predictions, we compared the frequency of colony extinction events between desert and savanna sites and evaluated whether the factors associated with colony extinction differed predictably across these habitat types.

## METHODS

The data herein were collected as part of a larger study examining site-specific selection on collective behavior and leader-follower interactions in social spiders (Pruitt et al. 2018). Pruitt et al. (2018) evaluated for and confirmed the presence of site-specific selection for social susceptibility at arid sites, using the survivorship data presented in this paper along with behavioral data not used here. The following data on correlates of extinction and risk of various selective agents, which are the unique focus of this paper, have not been published previously. The current paper therefore aims to identify differences in environmental selective forces that may underlie the site-specific selection described in Pruitt et al. (2018). There are no redundant conclusions between this paper and any other work published by our group.

All applicable institutional and/or national guidelines for the care and use of animals were followed

### Collection

We collected whole colonies of *S. dumicola* along roadside fences and *Acacia mellifera* trees at eight study sites. Sites were distributed across two precipitation transects: one extending north from the Namib Desert towards Angola (810 km), the other extending east from the Kalahari Desert towards Lesotho (981 km). Each transect contained four sites, two arid desert sites (Kalahari gradient: Upington [-28.403361, 21.071249] and Boegoeberg [-29.037819, 22.027999]; Namib gradient: Rehoboth [-23.209881, 17.092] and Kalkrand [-24.065027, 17.580452]) and two wetter savanna sites (Kalahari gradient: Ladysmith [-38.65655, 29.625249] and Weenen [-28.856239, 30.142306]; Namib gradient: Rundu [-18.299209, 19.407636] and Outjo[-20.233099, 16.354468]). In all, we collected 211 source colonies with populations of mature females spanning from 75-512 individuals per colony.

We collected colonies by placing a cloth pillowcase over the entire nest, then either prying the nest loose from fence wires or using garden clippers to separate the retreat from its substrate. We transported colonies in a climate-controlled vehicle to nearby hotels for separation/isolation. We individually dissected colonies by hand, separating each mature female spider into a 59ml deli cup for a 24-hour isolation period prior to assembling experimental colonies. When creating experimental groups, care was taken not mix individuals from multiple source colonies. In social *Stegodyphus*, variation in relatedness and/or familiarity has been demonstrated to alter colony behavior and performance (Modlmeier et al. 2014, Laskowski et al. 2016).

### Experimental Colony Assembly

We created a total of 242 experimental colonies, each containing 20 mature female spiders. Experimental colonies were housed in 390mL plastic cups, each containing three *A. mellifera* twigs to facilitate web construction. Several holes were punched into the bottom of each cup to allow for water drainage for instances of rain. Colonies were housed in their group cups for a total of 5 days at a temperature of 22-25oC before deployment into the field at their respective sites. During this time, colonies were able to acclimate to their new housing as well as build a retreat structure to disincentivize dispersal from their deployment location.

### Experimental Colony Deployment

We deployed our experimental colonies across the eight sites (28-33 colonies per site) using clothespins to attach individual cups to *A. mellifera* trees. Each colony was attached to its own tree to minimize interaction between colonies. Stegodyphus dumicola individuals will occasionally disperse to the nearest branch tip following deployment, but most frequently begin building their capture web directly atop the clothespin fixing their cup to the substrate. The aforementioned pre-deployment construction period aims to further suppress post-deployment dispersion and has been successfully implemented in prior field experiments using experimental colonies of S. dumicola (Grinsted et al. 2013, Keiser et al. 2014, Keiser et al. 2015, Wright et al. 2017). Colonies were not deployed onto trees that were actively being patrolled by predatory ants (genus *Anoplolepis* and *Crematogaster)*, as these ants have previously been demonstrated to destroy colonies before they can become established (Keiser et al 2015) and actively rove vegetation in search of prey (Doering et al. 2018). Colonies were deployed in an order that did not conflate one climate type with time of release: 1: Wet (Rundu, Nov 2016), 2: Wet (Outjo Nov 2016), 3: Dry (Rehoboth Nov 2016), 4: Dry (Kalkrand Nov 2016), 5: Dry (Upington Dec 2016), 6: Dry (Boegoeberg Dec 2016), 7: Wet (Ladysmith Dec 2016), 8: Wet (Weenen Jan 2017). Colonies were deployed at the same site from which their source colonies were collected.

### Colony Checks & Correlates of Extinction

To determine colony survival, colonies were left in their environments for the next two months. We then returned to each colony and determined whether it had survived or had gone extinct. We deemed a colony to have gone extinct if no living members of the society persisted.

To determine the potential causes of colony extinction, in addition to evaluating survival, we noted cues as to whether both surviving and extinct colonies had experienced attack by predatory ants or fungal outbreak. Substantial quantities of *Anoplolepis spp.* ant carcasses in the capture web, nest, and deployment container provided evidence of an attack by ants (Yip & Rayor 2011, Keiser et al 2015). *Stedodyphus dumicola* do not generally consume *Anoplolepis* as prey, and instead respond in an antipredator fashion by weaving hyperadhesive cribellate silk to stave ants off from the retreat (Henschell 1998, Keiser et al. 2015). Evidence of a fungal outbreak was noted if fungus was sprouting from carcasses of colony members, prey items, and the web (Henschel 1998). To measure the amount of prey captured by the colony we recorded the number of desiccated prey carcasses within each capture web and nest. Unlike many spider species that discard prey carcasses from their web after consumption, *S. dumicola* instead weave spent prey items directly into the three-dimensional retreat structure of their web. As such, an observer may count the number of prey items captured by the colony.

### Statistical Methods

We first established that our arid and wet sites experienced quantitatively different rates of precipitation by performing a t-test on each site’s monthly rainfalls over the five years preceding the experiment (2013-2017). We grouped Kalkrand (Namibia), Rehoboth (Namibia), Upington (S. Africa), and Boegoeberg (S. Africa) as our arid sites, and grouped Rundu (Namibia), Outjo (Namibia), Ladysmith (S. Africa), and Weenen (S. Africa) as our wet sites. Rainfall data specific to these sites was obtained from World Weather Online.

We used GLMMs to test for associations between colony persistence, factors reasoned to cause colony extinctions in *S. dumicola* (ant attacks, fungal outbreak, # prey captured), and habitat type (arid/wet) in R (ver. 1.1.383) using the package lme4 (Bates et al. 2014). We first compared overall rates of colony extinction between arid and wet sites by constructing a model with site type (arid/wet) as a fixed effect, site ID as a random effect, and colony survival (1/0) as a binary response variable. We used a binomial error distribution with a logit-link function.

To investigate how site type affected prey capture rates, we constructed a model including site type (arid/wet) as a fixed effect, site ID as a random effect, and the number of prey recovered from the colony’s web as a response variable. We fitted a Poisson error distribution and log-link function. This model included data only from colonies that survived the duration of our study, as colonies that perished would have had different and unknown lengths of time in which to capture prey.

To test for an association between mortality due to fungal outbreak and site type, we constructed a model with site type as a fixed effect, and site ID as a random effect, and incidence of fungal outbreak (1/0) as a binary response variable. We designated a binomial error distribution with a logit-link function. To focus on mortality-specific incidences of fungal outbreak, this model included data only from colonies that perished.

To test whether ant attacks occurred more frequently at wet sites, we constructed a model with site type as a fixed effect, site ID as a random effect, and evidence of ant attack (1/0) as a binary response. We designated a binomial error distribution with a logit-link function. We included both surviving and extinct colonies in this analysis; including both fatal and non-fatal attack events informs us of the overall frequency of predatory ant encounters. To evaluate whether colony *mortality* due to ant attack was more prevalent at wet sites, we constructed a model including site type as a fixed effect, site ID as a random effect, and evidence of ant attack as a binary response variable (1/0). We designated a binomial error distribution with a logit-link function. For this model, we used data only from colonies that had perished in order to focus on mortality-specific instances of these attacks.

In a *post hoc* analysis, we tested whether the number of prey present in a web was associated with fungal outbreaks. For this model we ran a GLMM with prey number, site type, and their interaction term as fixed effects, site ID as a random effect, and incidence of fungal outbreak as a binary response.

## RESULTS

As predicted, we found that our arid and wet site designations reflected qualitatively different precipitation regimes. Monthly precipitation at wet sites was found to be higher than at arid sites (*t* = 6.12 p <0.0001). Notably, wet sites experienced over double (~45mm per month) the rainfall of arid sites (~22mm per month) during their rainy season of December through March.

We found that colonies at arid and wet sites experienced comparable survival rates (~55% arid vs. ~44% wet), with a non-significant trend towards colonies surviving slightly better at arid sites (*site type*: z-stat = −1.787, df.resid= 239, p = 0.0739).

Among surviving groups that had not been raided by ants, colonies at wet sites had higher prey capture rates than colonies at dry sites (*site type*: z-stat = 8.853, df.resid = 94, p <0.0001, Fig 1). Surviving colonies at wet sites contained an overage of 10.3 prey items, whereas colonies at arid sites contained an average of 5.2 prey items.

**Figure 1:**
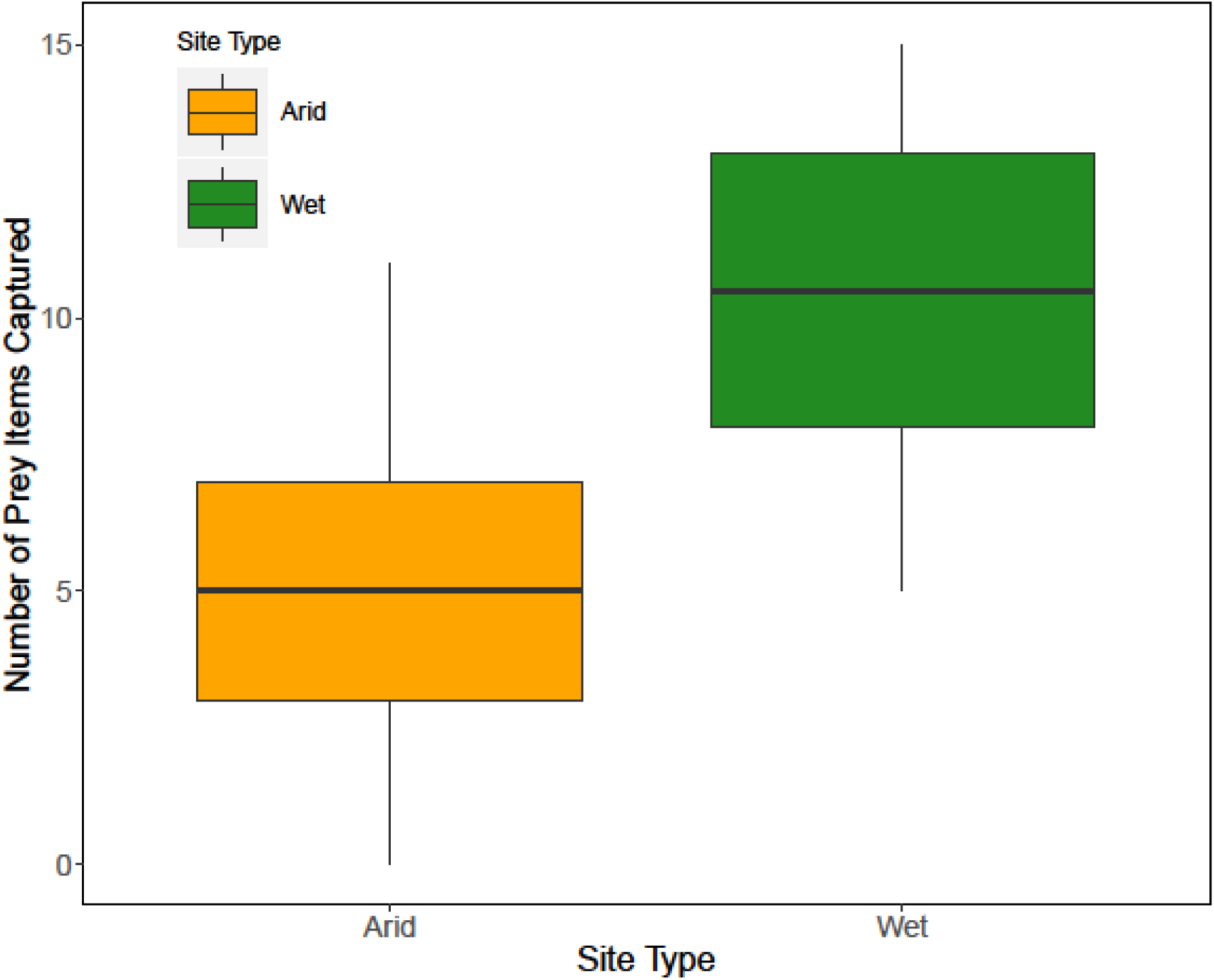
Number of prey items captured differs between site types. Number of prey items recovered from webs of colonies that survived the full duration of the study and did not display signs of ant attack in arid sites (orange) and wet sites (green). Boxes indicate the lower and upper quartiles; horizontal lines within boxes indicate the median, and whiskers extend to the 1.5 interquartile range from the box.

Our prediction that fungal outbreaks would be associated with colony extinctions at wet sites was corroborated by our data. A total of 31 colonies displayed evidence of fungal outbreak, with only 6 of those colonies surviving the duration of the study. Among colonies that perished, we detected a significant difference in the frequency of fungal outbreaks across site types (*site type*: z-stat = 3.248, df.resid = 120, p = 0.0012, Fig 2). Evidence of fungal outbreak was observed in 32% of the colonies that perished at wet sites and only 4% of colonies that perished at arid sites.

**Figure 2:**
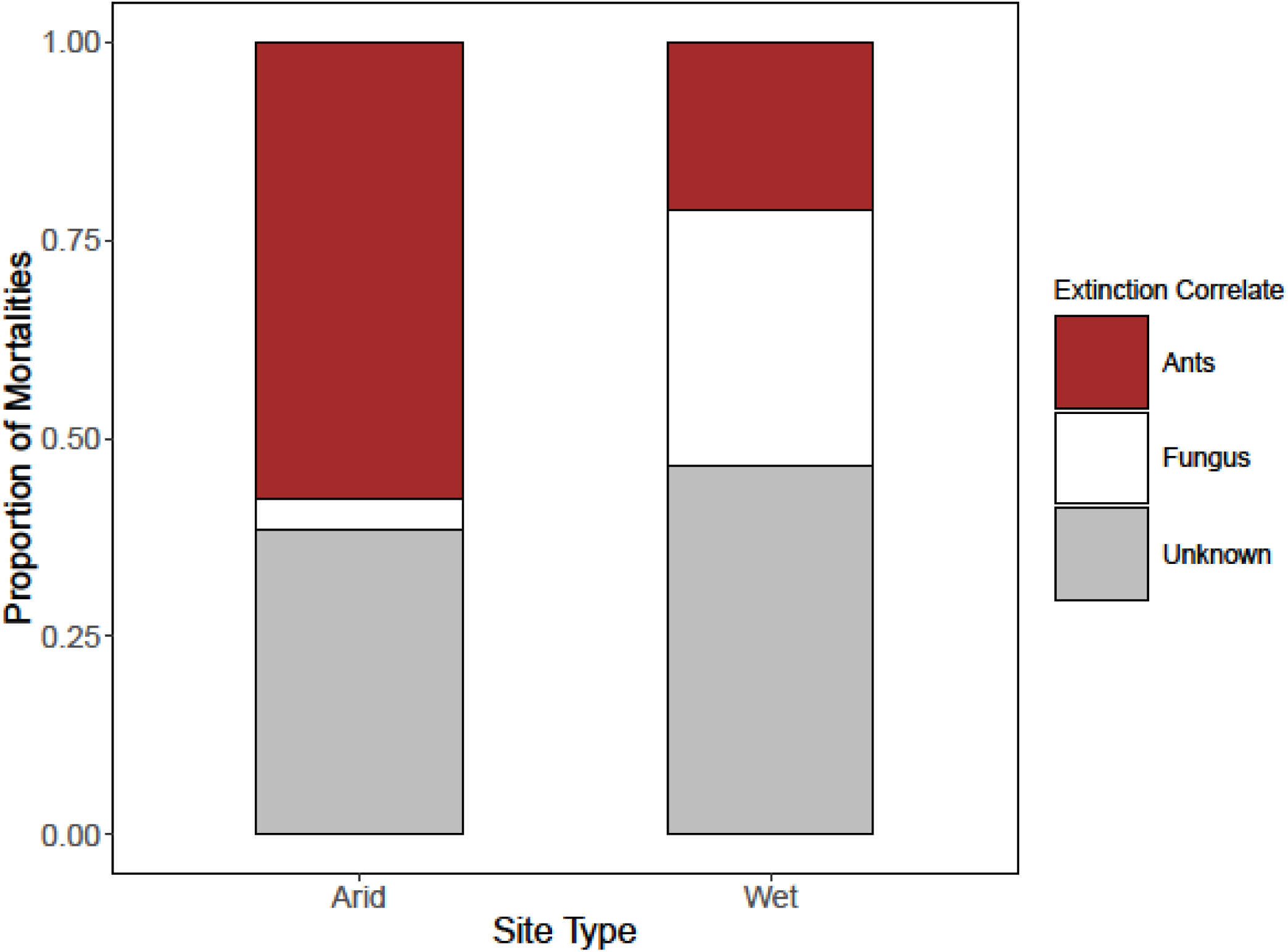
Correlates of Extinction Vary by Site Type. Proportion of total colony mortalities associated with ant attacks (red), fungal outbreaks (white), and unknown causes (grey) at arid (left) and wet (right) sites. Colonies containing *Anoplolepis spp*. carcasses were noted to be attacked by predatory ants, while colonies containing puffs of white fungus riddled through the nest were recorded as associated with fungal outbreaks.

We found that prey capture rates were positively associated with fungus across *both* site types, as the fixed effect of prey number was significant (*prey number*: z-stat = 2.367, Df.resid = 237, p = 0.01795, Fig 3) but the interaction term was not (*site type × prey number*: p = 0.0957). Therefore, colonies that had a greater number of prey carcasses in the web were more likely to experience fungal outbreaks, regardless of habitat type (Fig 3).

**Figure 3:**
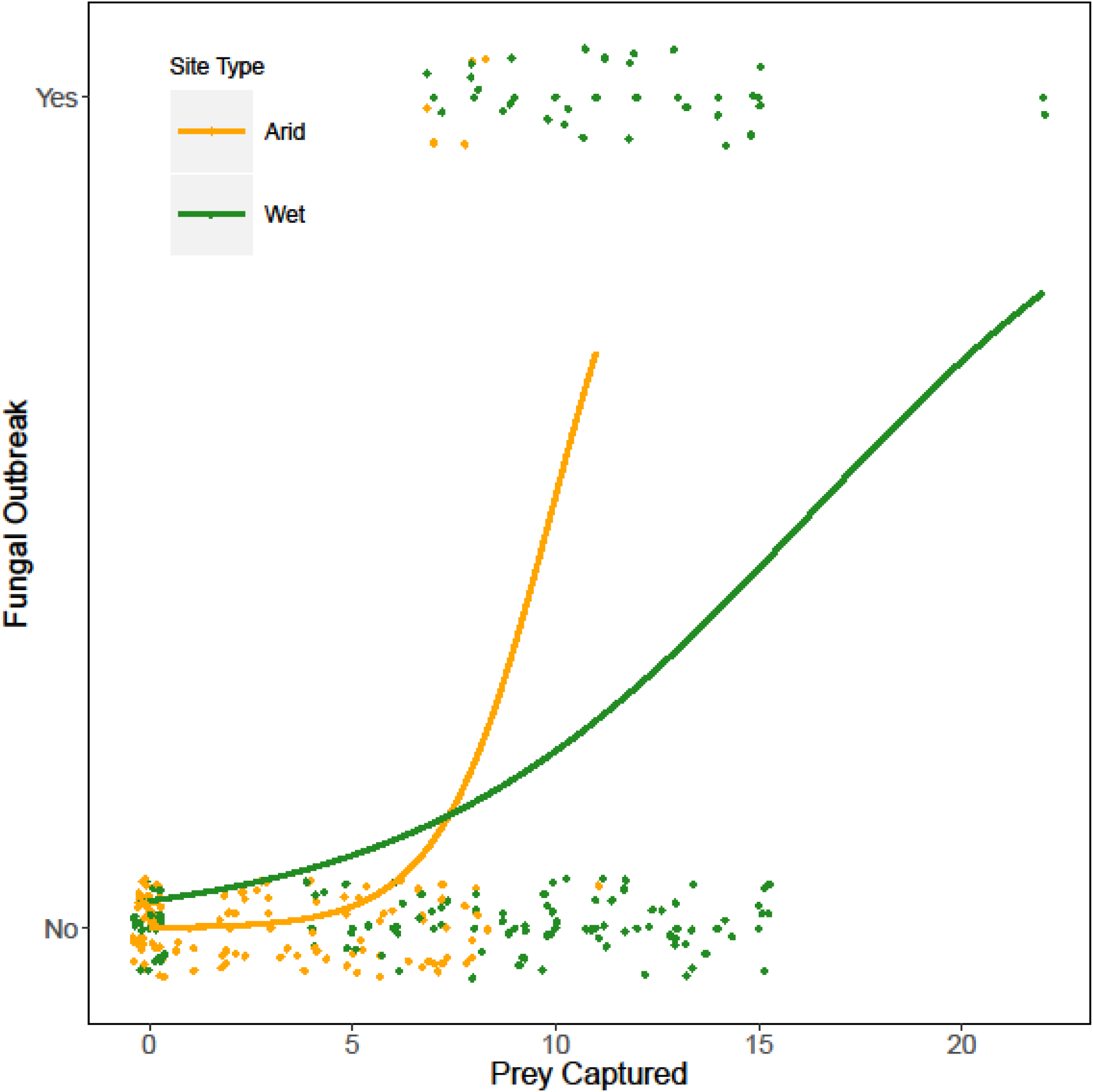
Incidence of Fungal Outbreak Versus Prey Capture Rate. Relationship between (binary) occurrence of fungal outbreak and prey capture rate at arid (orange) and wet (green) sites A logistic regression of predicted values for fungal outbreak as a function of prey capture is displayed by site type (arid: orange, wet: green). Points are jittered along the y axis to improve visibility

Our prediction that frequency of ant attack would be higher at wet sites was not corroborated by our data. Instead, the frequency of ant attack (including all attack events, survived by the colony or not) was higher in arid habitats than in wet habitats (*site type:* z-stat = −3.630, df.resid = 239, p = 0.0003). A total of 67 colonies had evidence of ant attack, 45 of which went extinct. Among colonies that perished, we found that the arid site type was associated with evidence of ant attacks (*site type*: z-stat = −4.027, df.resid= 120, p < 0.0001, Fig. 2). At arid sites, 58% of colony extinction events were associated with evidence of attacks by ants. In contrast, at wet sites only 21% of colony mortality events were associated with evidence of an attack by ants.

Of the sixty-seven colonies that displayed signs of an ant attack, only two colonies had *any* prey items remaining in their web. Both prey-containing webs yielded only one prey item each: the elytra of a very large beetle. This result is consistent with prior field observations that *Anoplolepis* ants will consume *S. dumicola*, nest webbing, and remnant prey carcasses during attacks.

## DISCUSSION

Identifying the forces driving group collapse and disbandment is key for understanding social evolution in many kinds of animals (Wray et al. 2011, Gordon 2013, Pruitt & Goodnight 2014). We provide evidence here that the likely drivers of colony extinction events differ across sites with contrasting precipitation regimes. We found that ant attacks were the most common source of colony extinction events at arid sites, whereas fungal outbreaks were most commonly associated with colony extinction events at wetter savannas. These differences may therefore impose divergent selection pressures on colony attributes at different site types, as has previously been shown in *S. dumicola* (Pruitt et al. 2018) and at least one other transitionally social species of spider (Pruitt & Goodnight 2014).

Our first prediction posited that colonies in wet sites would capture more prey than colonies in arid sites. When examining only surviving colonies, which had equal amounts of time to capture prey, we found that colonies in wet sites indeed had capture rates approximately twice those of colonies at arid sites. This is consistent with data demonstrating that savannas are more productive habitats than tropical deserts (Lieth 1973) and support larger numbers of phytophagous insects (Janzen and Schoener, 1968), which constitute the majority of prey for *S. dumicola* (Majer et al. 2018).

Consistent with predictions, fungal outbreaks were both more frequent and more commonly associated with extinction in wet sites than arid sites. One of the likely causes of these outcomes is that entomopathogenic fungus may require more humid conditions to persist and reproduce. This humidity may interact with the higher number of prey items in webs at wet sites to increase the chance of unconsumed prey spoiling and sprouting fungus. Wet sites may also have greater prey diversity with the potential to harbor and transmit fungus to *S. dumicola* colonies. Colonies at wet sites have a higher interaction rate with such prey, as evidenced by their incorporating a larger number of prey carcasses into their webs. Each of these factors has the potential to contribute to increased fungal outbreaks observed at wet sites. Recent metagenomic data from the webs, nests, and cuticles of *S. dumicola* confirm that microbial transmission from prey to spiders is common (Keiser et al. *in press*). Furthermore, prior observations of *S. dumicola* colonies infected by fungus note that individuals ‘became lethargic and died’ (Henschel 1998), indicating that fungal infection can lead to the mortality of individuals. However, it is also possible that fungal outbreaks may occur after a colony’s demise. This would give the appearance that a colony had been killed by a fungal outbreak. However, the fact that several colonies were observed to be infected by fungus and still functional lends support to the hypothesis that fungal outbreak precedes colony extinction. Further, were fungal outbreaks merely a byproduct of extinct colonies rotting in a wetter environment, the relationship between increased prey capture and fungal outbreak might have been weaker or nonexistent altogether.

At odds with our primary predictions, ant attacks appeared to be both more frequent at arid sites and more likely to lead to mortality for arid-dwelling colonies. Whether ants that prey upon *S. dumicola* are indeed more abundant at arid sites is yet unclear. We reason that an identical number of predatory ants at arid sites could prove more lethal to social spiders than the same number of ants at wet sites. This is because arid deserts, being less structurally complex than wetter savannas, are potentially easier for predatory ants to navigate and explore, leaving little opportunity for social spider colonies to go unnoticed. It is also likely that there are fewer alternative prey at arid sites that might distract predatory ants or satiate them. Deserts are known to limit the time and number of days that ant colonies can forage (Gordon 1991, Pinter-Wollman et al. 2012, Gordon 2013) and their prey preferences (Traniello et al. 1984), due to worker desiccation risk. If this is true in *Anoplolepis*, then this imposition is insufficient to decrease ant-driven lethality to social spider colonies at these sites.

We found that the majority of spider colonies attacked by ants had zero prey items in their web. Of the 67 colonies that displayed signs of ant attack, only 2 colonies had prey in their capture webs. Observations of interactions between *Anoplolepis* ants and *S. dumicola* note that attacking ants eat nearly everything in the capture web during their attack (Henschel 1998, Keiser et al. 2015). Additionally, should a colony survive an ant attack (as in 22 of 67 observed attacks in this study), a history of ant attacks is known to decrease colony foraging aggressiveness (Wright et al. 2017). This is costly, as collective foraging aggressiveness is associated with colony mass gain and colony persistence (Pinter-Wollman et al. 2017, Pruitt et al. 2018). Thus even non-lethal attacks by ants, if suffered early enough in a colony’s history, could render the group less likely to capture prey later.

Identifying spatial (between environments) and temporal (between seasons and across years; as in Bengston 2018) variation in selection pressures on social organisms helps understand the evolution of between-group differences in collective phenotypes. Such differences have been observed in many kinds of social animals (Wray et al. 2011, Scharf et al. 2012, Bengston 2014, Jandt et al. 2014, Jolles et al. 2018, Kamath et al. 2018). In particular, Pruitt et al. (2018) demonstrated the existence of site-specific selection on social susceptibility in *S. dumicola* at the same sites in which we evaluated drivers of extinction here. Colonies at arid sites exhibit a collective susceptibility to bold-phenotype individuals, increasing the colony’s collective aggression response in foraging for prey. Increased aggression at these arid sites, in turn, predicted colony survival. Colonies at wet sites showed no such susceptibility and demonstrated no change in collective foraging aggression in the presence of a bold-phenotype individual. Further, there was no link between colony survivorship and collective aggression at wet sites. Here we have begun to explore potential mechanisms underlying this contrasting pattern of selection by showing that social organisms in different habitat types appear to experience different environmental pressures, in the form of group extinctions. The correlates of colony extinction differed characteristically across site types in *S. dumicola*, with ant attacks being associated with colony extinction primarily at arid sites and fungal outbreaks being associated with extinctions at wet sites. These differences, in turn, may help to explain the differences in selection pressures on colony behavior seen in *S. dumicola* (Pruitt et al. 2018). We reason that similar site-specificity of group extinction drivers could be common in other social taxa.

## Author contributions

JNP and NPW conceived the experiment. BLM, JLL, CMW, GTC, and JNP performed the experiment. BLM analyzed the data. BLM and JNP wrote the manuscript; other authors provided editorial input

## Acknowledgements

Funding was provided by NSF IOS grants 1352705 and 1455895 to JNP, 1456010 to N.P.W. and NIH grant GM115509 to N.P.W. and J.N.P. Special thanks are due to Ian Van Wert for keeping science cool.

